# Single Plaque Proteomics Reveals the Composition and Dynamics of the Amyloid Microenvironment in Alzheimer’s Disease

**DOI:** 10.64898/2026.02.02.703320

**Authors:** Mengqi Chu, Ju Wang, Jay M. Yarbro, Ping-Chung Chen, Him K. Shrestha, Huan Sun, Mingming Niu, Zhen Wang, Sarah Harvey, Zhiping Wu, Yingxue Fu, Zuo-Fei Yuan, Haiyan Tan, Anthony A. High, Aijun Zhang, Xusheng Wang, Meifen Lu, Heather Sheppard, Geidy E. Serrano, Thomas G. Beach, Gang Yu, Yun Jiao, Junmin Peng

## Abstract

Alzheimer’s disease (AD) is characterized by amyloid plaques that form complex microenvironments in the brain. However, the molecular composition of these plaques and their temporal regulation are not well defined. Here, we developed a sensitive workflow for quantitative proteomic profiling of single plaques using refined laser capture microdissection and data-independent acquisition mass spectrometry (LCM-DIA-MS). From >200 plaques and control regions in AD mouse models (5xFAD and APP-KI) and human brains, we quantified >7,000 proteins, revealing stage-dependent, cell-type-related remodeling of the amyloid proteome (amyloidome). Temporal profiling uncovered early immune and lysosomal activation followed by engagement of RNA processing and synaptic pathways. Cross-model and cross-species analyses determined a conserved amyloidome including APOE, MDK, PTN, and HTRA1, validated by co-localization in imaging analysis. Network analysis highlighted modules in lipid transport, vesicle organization, and autophagy. These findings establish amyloid plaques as conserved, dynamic multicellular hubs that link amyloid accumulation to downstream cellular events.

## Introduction

Alzheimer’s disease (AD) is a progressive neurodegenerative disorder and the leading cause of dementia^1^. It is defined by the aggregation of Aβ peptides in extracellular amyloid plaques and hyperphosphorylated tau in intracellular neurofibrillary tangles^2^. These aggregates promote reactive microglial and astrocytic responses, driving neuroinflammation, synaptic loss, and ultimately neuronal death^2–4^. The amyloid cascade hypothesis, which states that Aβ aggregation is an initiating event in AD pathology, is supported by evidence that plaque deposition begins decades before clinical symptoms begin^5^. However, the detailed mechanisms linking the presence of amyloid plaques to progressive neural dysfunction remain unclear.

AD progression is manifested by the cellular microenvironment surrounding amyloid plaques. While Aβ fibrils form the insoluble plaque core, the microenvironment undergoes dynamic changes. Microglia are rapidly recruited to sites of Aβ deposition^6^, where their phagocytic activity may initially be protective^7^. During disease progression, microglia shift to a chronically activated, dysfunctional state marked by impaired Aβ clearance and excessive release of pro-inflammatory cytokines, resulting in neuroinflammation^8^. This persistent neuroinflammation causes synaptic damage and neuronal toxicity. Astrocytes also become reactive, showing apoptotic and senescence phenotypes and functional dysfunction in calcium signaling and glutamate buffering^9^. Additionally, Aβ may induce membrane damage of nearby dystrophic neurites, driving phosphorylated tau accumulation^10^. These cellular and structural changes create a self-perpetuating cycle of inflammation and neurodegeneration around amyloid plaques.

Although microglia, astrocytes, and dystrophic neurites are known to cluster tightly around amyloid plaques^6,11^, the precise molecular composition of this multicellular niche and its alterations over time remain unknown. Bulk-tissue proteomics^12–15^ studies can identify numerous AD-associated proteins but average signals across cell types and regions, obscuring the molecular signatures unique to plaque microenvironment^16,17^. In spatial proteomics studies^18^, laser capture microdissection (LCM)-MS has yet to achieve single-plaque proteomic depth due to sensitivity constraints^17,19–22^. Antibody-based imaging also provides spatial context but remains limited in multiplexing and unbiased discovery capacity^16^.

Recent advances in data-independent acquisition mass spectrometry (DIA-MS) enable reproducible, high-coverage, and quantitative proteomic analysis by systematically fragmenting all ions within defined mass ranges, enabling sensitive protein analysis from minute protein samples^23,24^. Here, we present an ultrasensitive LCM-DIA-MS workflow that quantifies >7,000 proteins from individual amyloid plaques. Applying this approach to >200 individual plaques and matched non-plaque regions from 5xFAD^25^ and APP-KI mouse models^26^ at multiple ages, as well as postmortem human AD brains, we uncovered temporally ordered remodeling of the plaque proteome, defined a conserved cross-species amyloidome, and identified coordinated multicellular networks. These studies not only establish an advanced spatial proteomics platform, but also reveal amyloid plaques as dynamic, conserved hubs rather than inert deposits, providing a scalable framework for dissecting plaque heterogeneity and stage-specific pathogenesis in AD.

## Results

### High-resolution single-plaque proteomics by LCM-DIA-MS

To characterize the proteomic landscape of individual amyloid plaques, we developed a workflow for the proteomic profiling of single plaques from mouse and human brain tissue at single-plaque resolution (∼50 μm diameter on 12-µm-thick slides). We analyzed a total of 226 samples, isolated from two common AD mouse models (5xFAD and APP-KI) and human postmortem AD brains (**Fig. 1a, Tables S1, S2**), using LCM-DIA-MS (**Fig. 1b**). Briefly, brain tissues from mice and AD patients were embedded in Optimal Cutting Temperature (OCT) compound, sectioned, and stained with X-34 to visualize plaques. LCM was then used to isolate individual plaques, as well as adjacent non-plaque regions as controls (**Fig. 1c**).

**Fig. 1.**
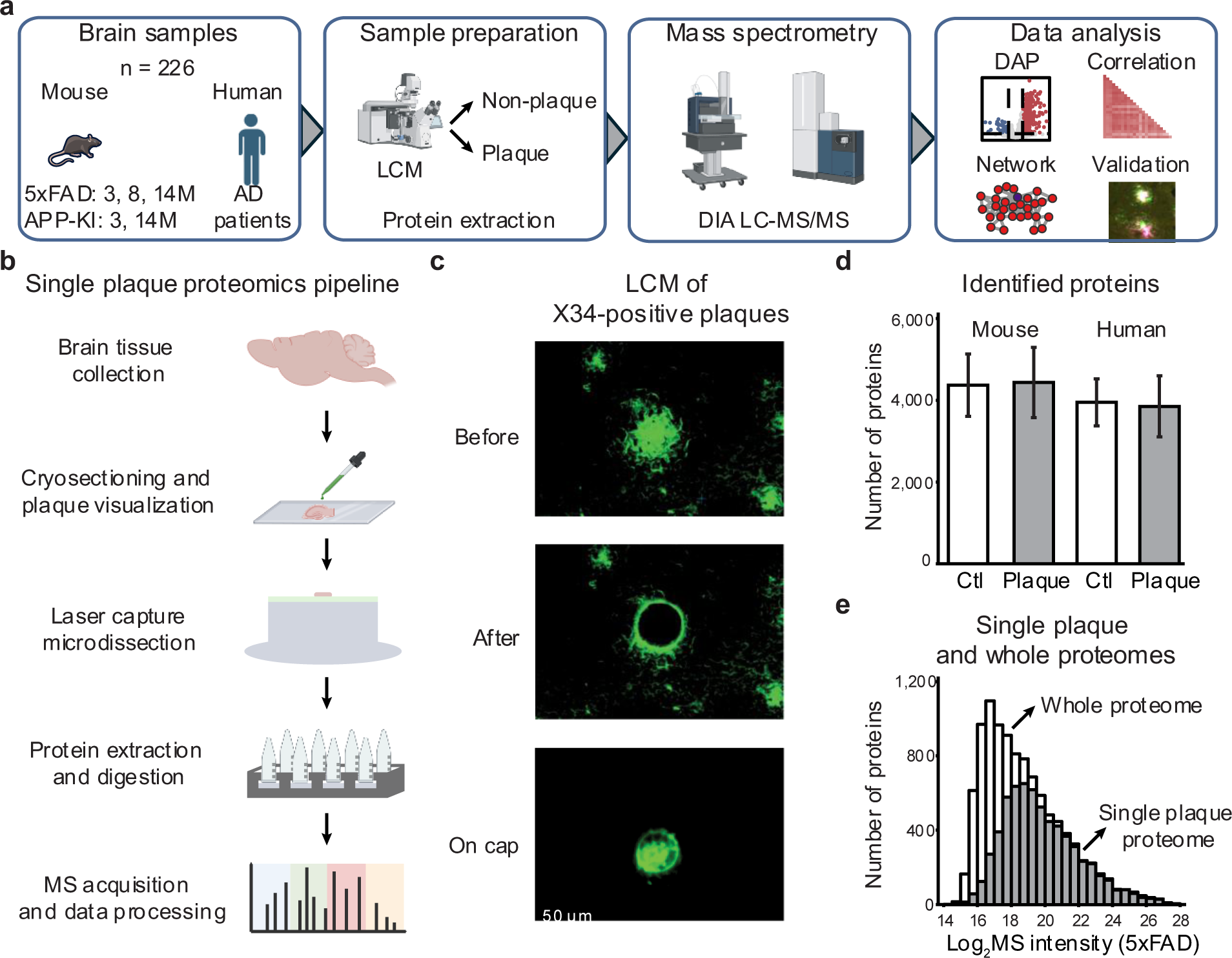
LCM-based single plaque proteomics study workflow shows comparable coverage to prior whole-proteome analyses. **a**, Study design: amyloid plaques isolated by laser capture microdissection (LCM) were analyzed by LCM-DIA-MS (5xFAD, APP-KI, human AD; *n* = 226). **b**, Workflow for quantifying plaque and non-plaque proteomes. **c**, Example images of an amyloid plaque before and after LCM. **d**, Quantified proteins in plaque vs. non-plaque regions across species. **e**, Comparison of quantified proteins from prior whole-proteome vs. current single-plaque analysis.

While the conventional proteomics workflow enables profiling of pooled samples (e.g., at least 500 plaques)^17^, it is not optimized for nanogram-scale inputs (∼2 ng per plaque)^19^, when sample loss and instrument sensitivity become limiting factors^27^. To address this, we developed a low-input LCM-DIA-MS protocol. LCM was used to isolate individual plaques, with protein extraction and digestion carried out directly on the LCM cap to minimize transfer loss. For this step, 0.02% *n*-dodecyl-β-D-maltoside (DDM) was added to facilitate protein extraction and reduce peptide adsorption^28,29^. To further reduce loss, the desalting step typically after trypsin digestion was omitted. Finally, we employed a narrow C18 column (50 µm inner diameter) to replace the standard 75 µm column for LC-MS/MS on a timsTOF SCP instrument using optimized settings^30^.

To evaluate protein contaminants in this workflow, we processed five negative controls by placing empty caps onto mouse brain tissue without LCM capturing (**Extended Data Fig. 1a**). At a 1% false discovery rate (FDR) using the DIA-NN database search program^31^, ∼500 proteins were detected in control samples, whereas >4,000 proteins were consistently identified from single-plaque samples (**Extended Data Fig. 1b**). Many proteins detected in controls overlapped with the most abundant proteins in plaques, indicating background binding during cap-tissue contact, but clear distinction between the controls and plaques supports that identified proteins originate primarily from plaque tissue (**Extended Data Fig. 1c**). Further, by analyzing 1, 3, and 10 tissue captures, we observed proportional increases in precursor and protein identifications (**Extended Data Fig. 1d-f**), demonstrating that signals derive from targeted captures rather than background contamination.

To evaluate proteome depth, we compared our single-plaque dataset to a deep whole-proteome reference from 5xFAD mouse brain^17^. On average, >4,000 proteins were identified from both individual plaque and non-plaque regions in human and mouse tissues (**Fig. 1d**). A total of 7,048 proteins were detected from all LCM samples, in which 6,424 overlap with the whole-brain proteome (10,369 proteins). The overlapping proteins included nearly all medium- to high-abundance species, demonstrating that our workflow captures the major expressed proteome from single plaque samples (**Fig. 1e**).

Thus, this LCM-DIA-MS workflow enables highly sensitive, reproducible, and quantitative profiling of individual amyloid plaques from low inputs of brain tissue. It provides a scalable platform to determine proteomic changes across disease stages and species.

### Temporal proteome remodeling in 5xFAD plaques

To gain deeper insight into amyloid plaque development in AD, we applied the LCM-DIA-MS workflow to 5xFAD mice at three ages (3-, 8-, and 14-months). A total of 123 dissected regions, including plaques and adjacent non-plaque areas, were profiled, with ∼20 replicates per group. Across all samples, 7,048 unique proteins (∼4,500 proteins per sample) were identified at a 1% FDR (**Fig. 2a, Extended Data Fig. 2a, Table S3**). Following protein quantification (**Extended Data Fig. 2b**), the abundance of the Aβ-derived tryptic peptide LVFFAEDVGSNK was highly enriched in plaques compared to non-plaque regions and showed a significant age-dependent increase (**Fig. 2b, Extended Data Fig. 2c**), confirming its role as a positive control for AD amyloid plaques. Principal component analysis (PCA) clearly separated plaque and non-plaque proteomes at each disease stage (**Extended Data Fig. 2d**).

**Fig. 2.**
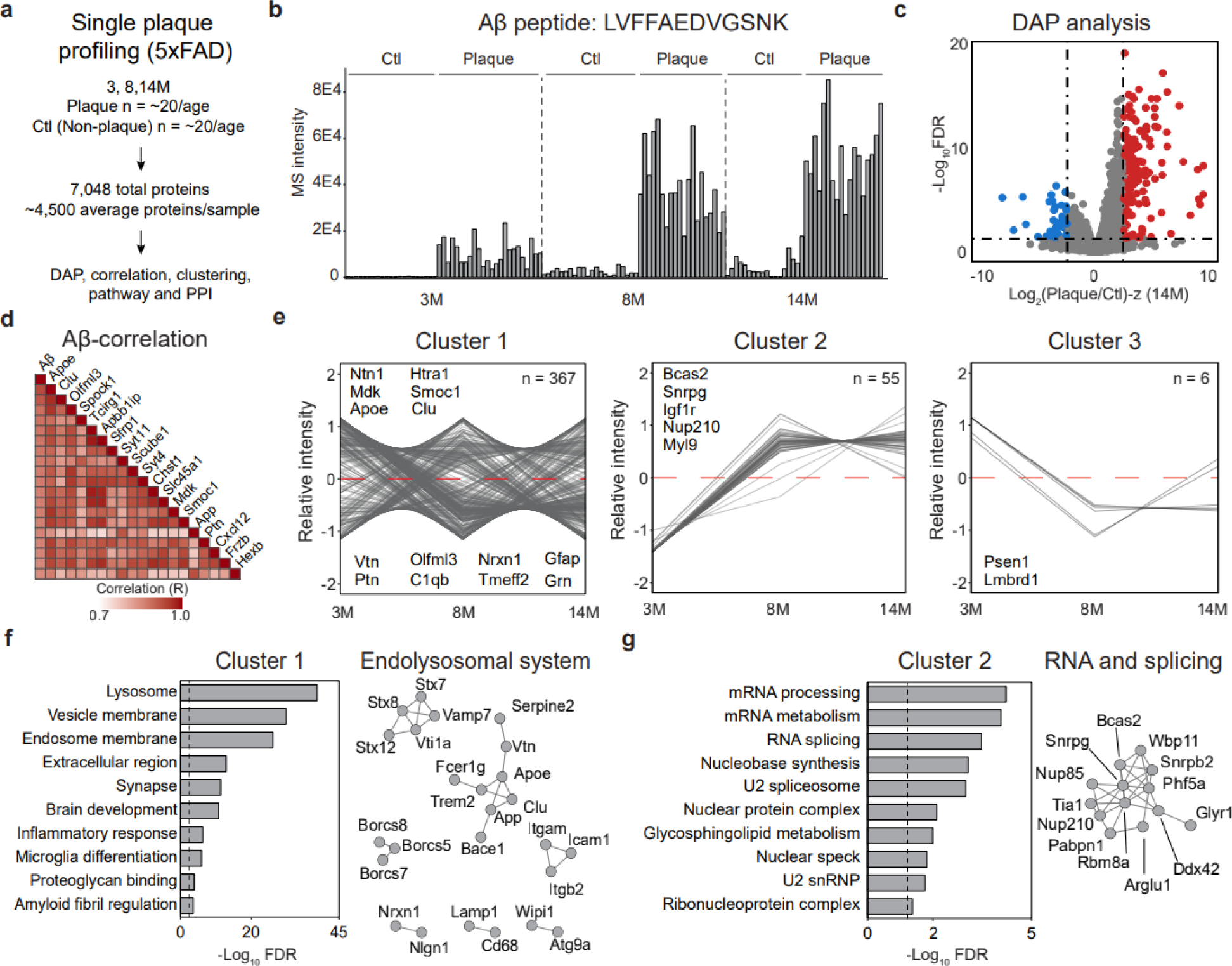
Proteome profiling in 5xFAD identifies temporal clusters and pathway-specific alterations in amyloid plaques. **a**, Paired plaque and non-plaque proteome profiling (*n* = 123) quantified 7,048 proteins. **b**, MS-based quantification of 5xFAD Aβ using peptide LVFFAEDVGSNK. **c**, Volcano plot of proteins enriched in plaque vs. non-plaque regions. **d**, Top Aβ-correlated proteins in the 5xFAD amyloid plaque proteome. **e**, Temporal plaque clusters: early/constant aggregation (cluster 1), late plaque accumulation (cluster 2), and decline over time (cluster 3). **f**, Pathway analysis of cluster 1 highlights endolysosomal and immune functions, and PPI network of endolysosomal DAPs. Dashed line represents cutoff for significance. **g**, Pathway analysis of cluster 2 highlights splicing and mRNA processes, and PPI network of splicing- and RNA-related DAPs. Dashed line represents cutoff for significance.

To identify amyloid plaque-associated proteins, differential abundance analysis was performed for each age group using two cutoffs (FDR <0.05 and |log_2_FC-z|>2.5). This analysis revealed a progressive increase in the number of differentially abundant proteins (DAPs) with age: 188 DAPs at 3-months (129 enriched in plaques), 195 at 8-months (159 enriched in plaques), and 220 at 14-months (181 enriched in plaques), suggesting that proteomic differences between plaque and non-plaque regions increase during AD progression (**Fig. 2c, Extended Data Fig. 3a, Table S4**). In total, 431 unique DAPs were identified across all ages, with 50 plaque-enriched DAPs at all stages including Aβ and Apoe, which were consistent with whole-proteome profiling of AD and control brain tissue^16^ (**Extended Data Fig. 3b, 3c**).

To investigate the molecular associations within amyloid plaques in the 5xFAD model, we performed a correlation analysis between Aβ and all other proteins. This analysis identified 57 proteins that were highly positively correlated with Aβ (R > 0.8). Among these, the top 20 proteins (**Fig. 2d)** included 11 previously reported Aβ-associated proteins such as Apoe, Clu, and Mdk^14^. To further characterize the proteomic dynamics, DAPs were grouped based on their fold-change patterns across ages, revealing three distinct temporal cluster patterns (**Fig. 2e, Table S5**). The patterns included: cluster 1 (early/constant aggregation, 367 proteins), encompassing most DAPs such as Ntn1, Mdk, Smoc1, Clu, and Htra1; cluster 2 (late accumulation, 55 proteins), including proteins such as Vim, Spock2, and Igf1r; and cluster 3 (decline over time, 6 proteins).

To investigate the biological functions of the temporal clusters, pathway analysis was performed for each (FDR < 0.05, **Table S6**). Cluster 1 was significantly associated with endolysosomal and immune functions, synaptic regulation, and amyloid fibril formation, indicating that these processes are engaged early in plaque development and remain active over time (**Fig. 2f**). In contrast, Cluster 2 was highly enriched for proteins involved in RNA processing and mRNA splicing, suggesting a key role for transcriptional and post-transcriptional regulation at later stages of disease progression (**Fig. 2g**).

Collectively, temporal proteome profiling of single plaques in the 5xFAD mouse model revealed dynamic molecular signatures associated with Aβ pathology. The three clusters highlight the roles of specific proteins and biological processes during distinct stages of amyloid plaque development.

### Cellular composition in the amyloid plaque microenvironment

To characterize the cellular components in the amyloid plaque microenvironment, we performed multiplexed immunofluorescence staining in 5xFAD mice (14-month-old, **Fig. 3a**). Amyloid plaques, astrocytes, microglia, and neurons were labeled with antibodies against Aβ, Gfap, Iba1, and NeuN/Map2, respectively (**Fig. 3b**). Within plaque-associated regions, partial colocalization of Aβ with distinct astrocytic, microglial, and neuronal markers was observed.

**Fig. 3.**
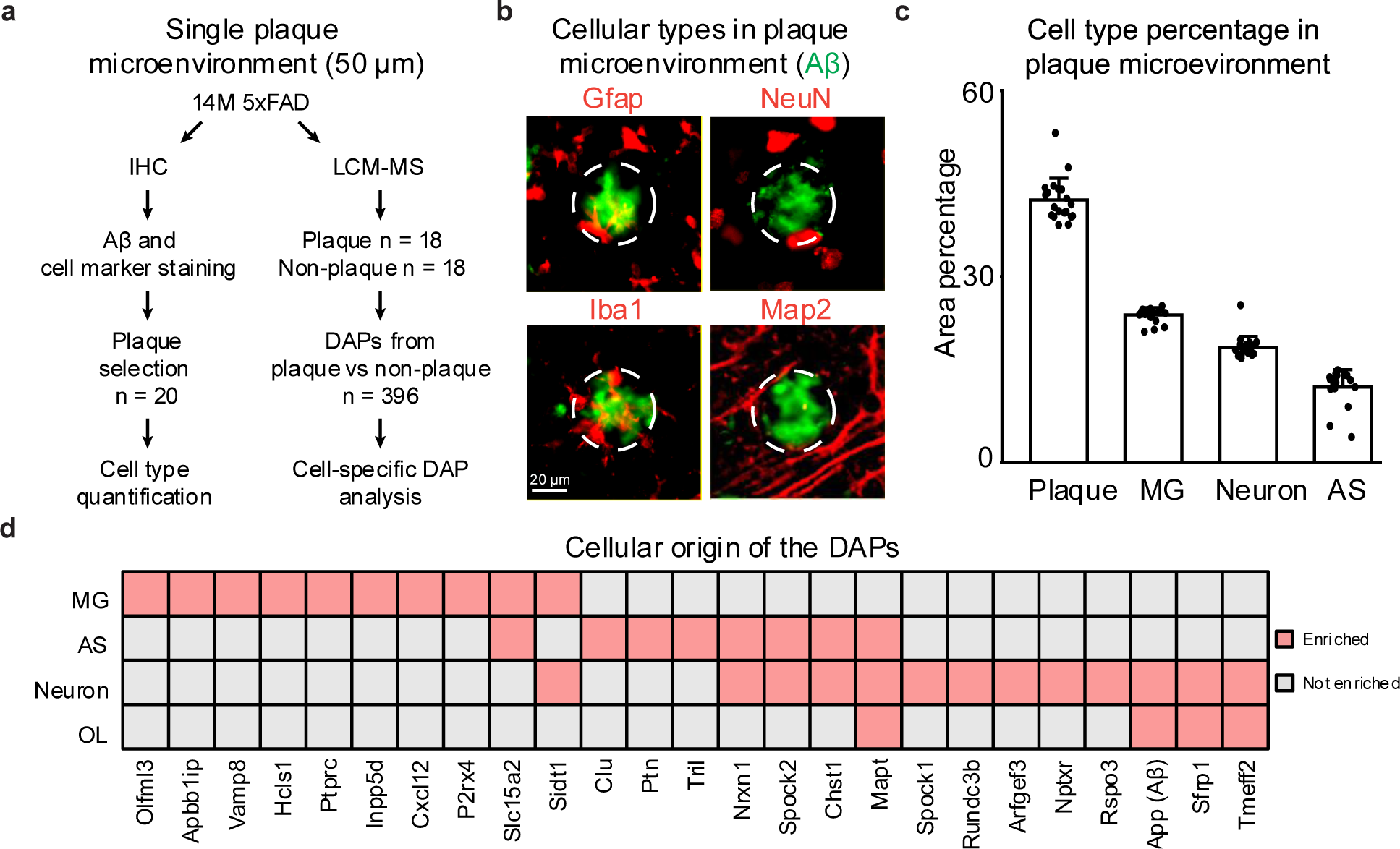
Cell type analysis of amyloid plaque microenvironment in 5xFAD. **a**, Workflow of single-plaque microenvironment analysis from 14M 5xFAD mice (n = 36 plaques). **b**, Immunofluorescence staining of amyloid plaques (green) and cell-type-specific markers (red) in the surrounding microenvironment. **c**, Relative proportions of major cell types within the amyloid plaque microenvironment (n = 20). **d**, Cellular origins of DAPs detected in plaque versus non-plaque proteomes.

To quantify cell-type composition within plaques, we analyzed the plaque microenvironments (50 μm diameter) for each co-staining combination of Aβ and different cell-type markers (**Extended Data Fig. 4a**). Using QuPath’s^32^ cell detection function, areas positive for Aβ and each cell marker were identified to calculate relative proportions (**Extended Data Fig. 4b, Table S7**). On average, Aβ occupied 42% of the staining area in plaque microenvironment, while microglia accounted for 24% and astrocytes for 12%. For neurons, the combined labeling areas of NeuN (staining neuronal soma) and Map2 (staining neuronal processes) represented 18% of the plaque niche. Overall, these data support that plaque microenvironments are predominantly occupied by the staining of Aβ and activated glial cells, with fewer neuronal components (**Fig. 3c**).

To further determine the origins of DAPs identified by LCM-MS, we traced the 396 DAPs detected in the 5xFAD plaques back to their cellular expression profiles, using a reference RNA-seq cell type dataset^33^. From 11,020 high-confidence cell-enriched transcripts, some DAPs were mapped to specific cell types, including 54 neuronal, 39 microglial, 18 astrocytic, and 18 oligodendrocytic proteins (**Table S8**). Notably, among the top 100 DAPs with the highest fold changes, 25 could be assigned, comprising 13 neuronal, 10 microglial, 8 astrocytic, and 4 oligodendrocytic specific proteins (**Fig. 3d**).

Overall, the plaques in 5xFAD mice exhibit a glia-dominated microenvironment in addition to Aβ aggregates, with microglia and astrocytes showing greater representation and proteomic enrichment than neurons.

### Conserved amyloid signatures and temporal dynamics in APP-KI plaques

To validate the proteomic signatures of amyloid pathology in the 5xFAD mice, we extended the single-plaque proteomic analysis to the humanized APP-KI mouse model without APP overexpression. In total, 75 LCM samples, including plaques and non-plaque controls from 3- and 14-month-old APP-KI mice, were processed using an identical LCM-DIA-MS workflow (**Fig. 4a, Tables S1, S2**). This analysis identified 6,701 proteins after stringent quality control, ensuring cross-model comparability (**Extended Data Fig. 5a, Table S9**). Following quantification and normalization (**Extended Data Fig. 5b**), the APP-KI-specific humanized Aβ peptide (LVFFA***G***DVGSNK) was highly enriched in plaques and increased with age (**Fig. 4b, Extended Data Fig. 5c**). Like 5xFAD, PCA distinguished plaque from non-plaque proteomes in the APP-KI model (**Extended Data Fig. 5d**).

**Fig. 4.**
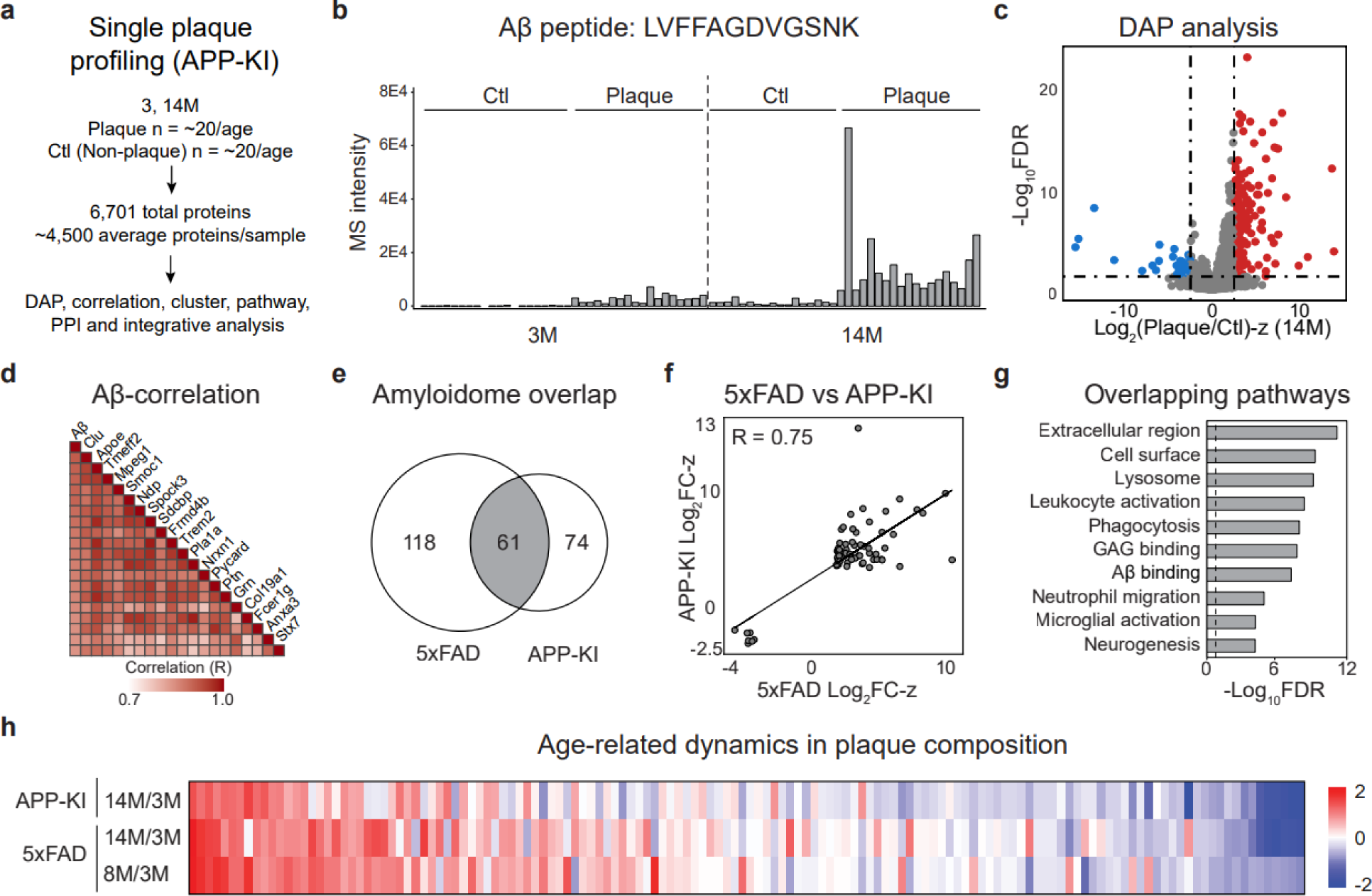
APP-KI plaque proteomics reveal shared amyloidome signatures, temporal clusters, and pathway alterations with 5xFAD. **a**, Paired plaque and non-plaque proteome profiling (*n* = 75) quantified 6,701 proteins. **b**, MS-based quantification of APP-KI Aβ using peptide LVFFAGDVGSNK. **c**, Volcano plot of proteins enriched in plaque vs. non-plaque regions. **d**, Top Aβ-correlated proteins in the APP-KI amyloid plaque proteome. **e**, Overlap of amyloidome DAPs in 5xFAD (14M) and APP-KI (14M). **f**, Correlation of consistent DAP Log_2_FC-Z values between 5xFAD and APP-KI. **g**, Pathway analysis of overlapping DAPs highlights extracellular, lysosomal, and immune proteins. Dashed line represents cutoff for significance. **h**, Heatmap illustrating temporal clustering across 5xFAD and APP-KI plaque progression.

Differential abundance analysis, using the same statistical thresholds as 5xFAD (FDR < 0.05, |log₂FC-z| > 2.5), identified 177 significantly altered proteins at 3-months and 165 at 14-months in APP-KI plaques (**Fig. 4c, Table S10**). Most DAPs were also increased in the plaque proteome. The correlation analysis identified 45 proteins that were highly positively correlated with Aβ (R > 0.8). Among these, the top 20 proteins included previously reported Aβ-associated proteins such as Apoe, Clu, and Ptn^14^ (**Fig. 4d**). Cross-model comparisons of the DAPs at each age are summarized in **Table S11**. At 14-month age, both models yielded the largest number of DAPs, with 100 overlapping proteins (**Fig. 4e**), and their correlations of log₂FC-z scores across models demonstrate significance (R = 0.85; **Fig. 4f**), further supporting the robustness of these shared signatures of amyloidosis.

Pathway analysis of APP-KI plaques revealed enrichment of immune-related processes, including leukocyte activation, phagocytosis, and microglial activation, which were also shared with 5xFAD (**Fig. 4g, Table S12**). Temporal analysis of the APP-KI and 5xFAD datasets identified 140 proteins exhibiting consistent regulation across models (**Fig. 4h**). These findings define a conserved amyloidome across two independent mouse models of amyloidosis, highlight shared age-dependent proteomic dynamics.

### Co-expression network modules and hub proteins in AD mouse plaques

To gain insight into the complex molecular networks in the plaque microenvironment, we performed co-expression network analysis using multiscale embedded gene co-expression network analysis (MEGENA)^34^. The entire proteomic datasets from 5xFAD and APP-KI models were combined, normalized, and used to construct a co-expression network (**Fig. 5a, Extended Data Fig. 6a**). The resulting network comprised 134 modules that passed the module significance threshold (p < 0.05; **Table S13, Extended Data Fig. 6b, c),** indicating well-structured modular organization.

**Fig. 5.**
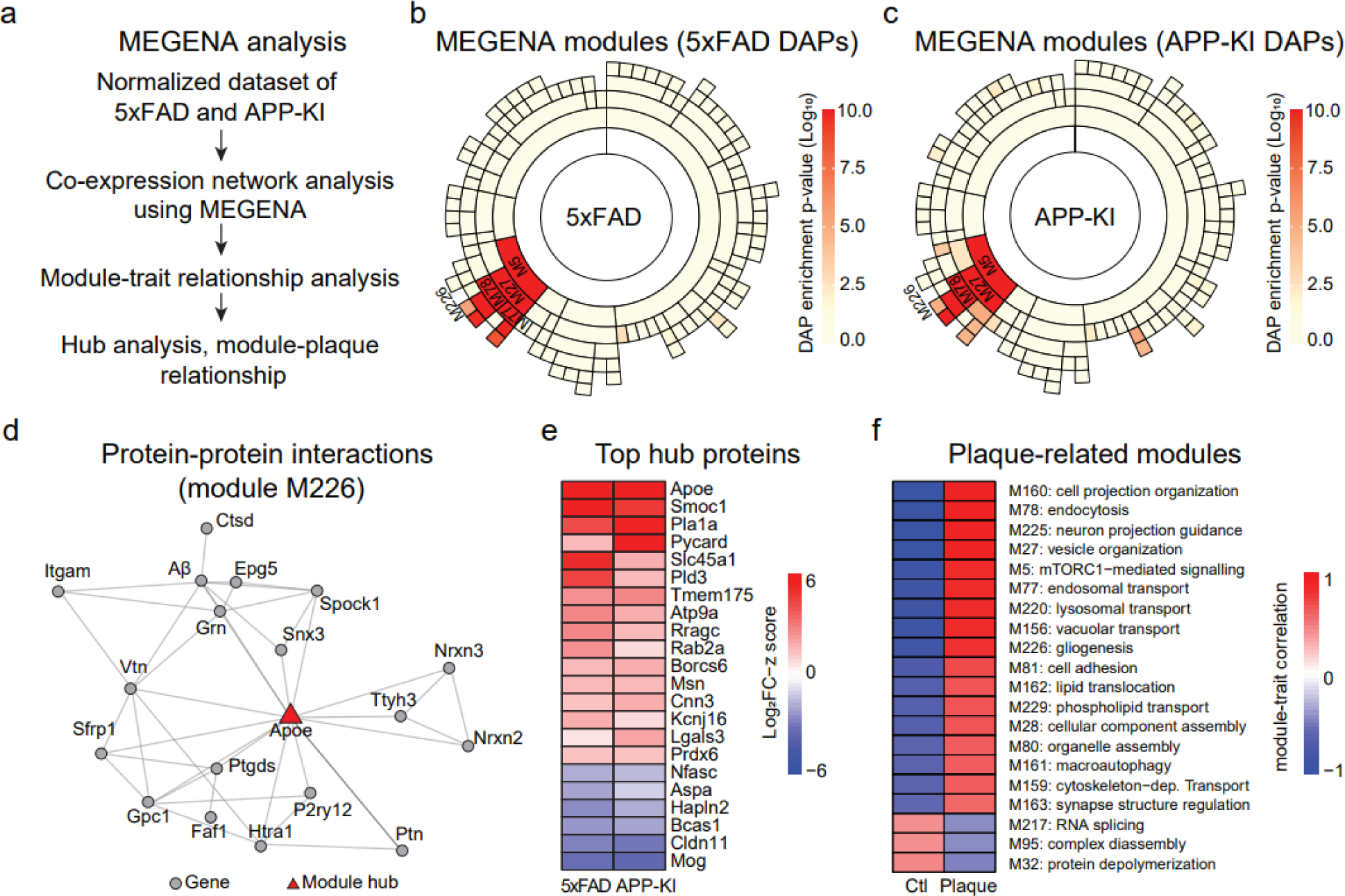
Co-expression network analysis identifies hub proteins and plaque-associated modules in 5xFAD and APP-KI mouse models. **a**, Schematic overview of the MEGENA workflow including network construction, module-trait relationship analysis, and hub protein identification. **b,c**, Sunburst plots showing module enrichment of DAPs in 5xFAD (b) and APP-KI (c) datasets, with significantly enriched modules highlighted in red gradient. **d**, Representative PPI network of module identified using M226, with genes represented as nodes and edges denoting experimentally supported interactions. Module hub is shown in red triangle. **e**, Heatmap of top hub proteins across 5xFAD and APP-KI models. Log_2_FC-z values from 5xFAD and APP-KI analysis were used for the heatmap. **f**, Modules correlating with plaque or non-plaque samples. Modules were qualitatively annotated using top enriched biological processes. Positive and negative correlations with plaque or non-plaque are indicated by red and blue, respectively.

To identify biological modules enriched for DAPs across AD mouse models, proteins were ranked based on the absolute value of their log_2_FC-z scores and tested for enrichment against the modules identified using 10,000 permutations. This analysis revealed 16 and 23 modules significantly enriched for DAPs in the 5xFAD and APP-KI models, respectively (**Fig. 5b, 5c**). Importantly, several top modules including M5 (macroautophagy), M27 (vesicle organization), M78 (endocytosis), and M226 (gliogenesis) were consistently enriched in both mouse models, suggesting shared biological responses associated with amyloid plaque pathology (**Extended Data Fig. 6d**). Module M226, which contained Aβ peptides, identified APOE as a key hub protein within its PPI subnetwork (**Fig. 5d**). Several additional proteins within this module, including Ptn and Htra1, were enriched in plaques and have been independently reported to co-localize with amyloid deposits in previous proteomic and immunohistochemistry studies^16,35^.

Hub proteins are highly connected nodes within molecular networks that act as central organizers and regulators of essential cellular processes^34^. We extracted MEGENA-derived hub proteins passing the FDR threshold, showing consistent trends between 5xFAD and APP-KI models, and exhibiting an absolute log_2_FC-z greater than 1.5. This analysis identified 22 hub proteins, including 16 upregulated and 6 downregulated proteins within the plaque regions (**Fig. 5e**). Among these, Apoe, Smoc1, Pycard, Pld3, and Tmem175 emerged as prominent upregulated hubs.

To determine modules associated with plaque versus non-plaque regions, we applied a module-trait correlation analysis in the MEGENA program. At an FDR threshold of 0.05 and an absolute correlation coefficient cutoff of 0.4, we identified 20 modules, of which 17 were positively correlated in the plaque region and 3 were positively correlated in the non-plaque region (**Fig. 5f**). In the plaques, M162 and M229 indicated upregulated lipid transport activity, and M161 and M78 suggested defective clearance and loss of cellular homeostasis. In the non-plaque area, M32 and M95 implicated deregulation of protein complex assembly. Thus, the MEGENA analysis identified several biologically relevant modules linked to amyloid pathogenesis and identified conserved hub proteins across models, providing molecular insight into the coordinated network architecture underlying plaque formation and progression.

### Human-mouse conserved amyloidome signatures

To validate amyloid-associated proteomic signatures, we extended single-plaque proteomic analysis to human AD cases. A total of 28 LCM samples, including plaques and adjacent non-plaque controls, were processed using the LCM-DIA-MS workflow (**Fig. 6a**). This yielded 5,865 proteins for comparison across species (**Extended Data Fig. 7a, 7b; Table S14**). As expected, the human Aβ peptide (LVFFAEDVGSNK) was markedly enriched in human plaques (**Fig. 6b, Extended Data Fig. 7c**). PCA also separated plaque and non-plaque proteomes in human cases (**Extended Data Fig. 7d**).

**Fig. 6.**
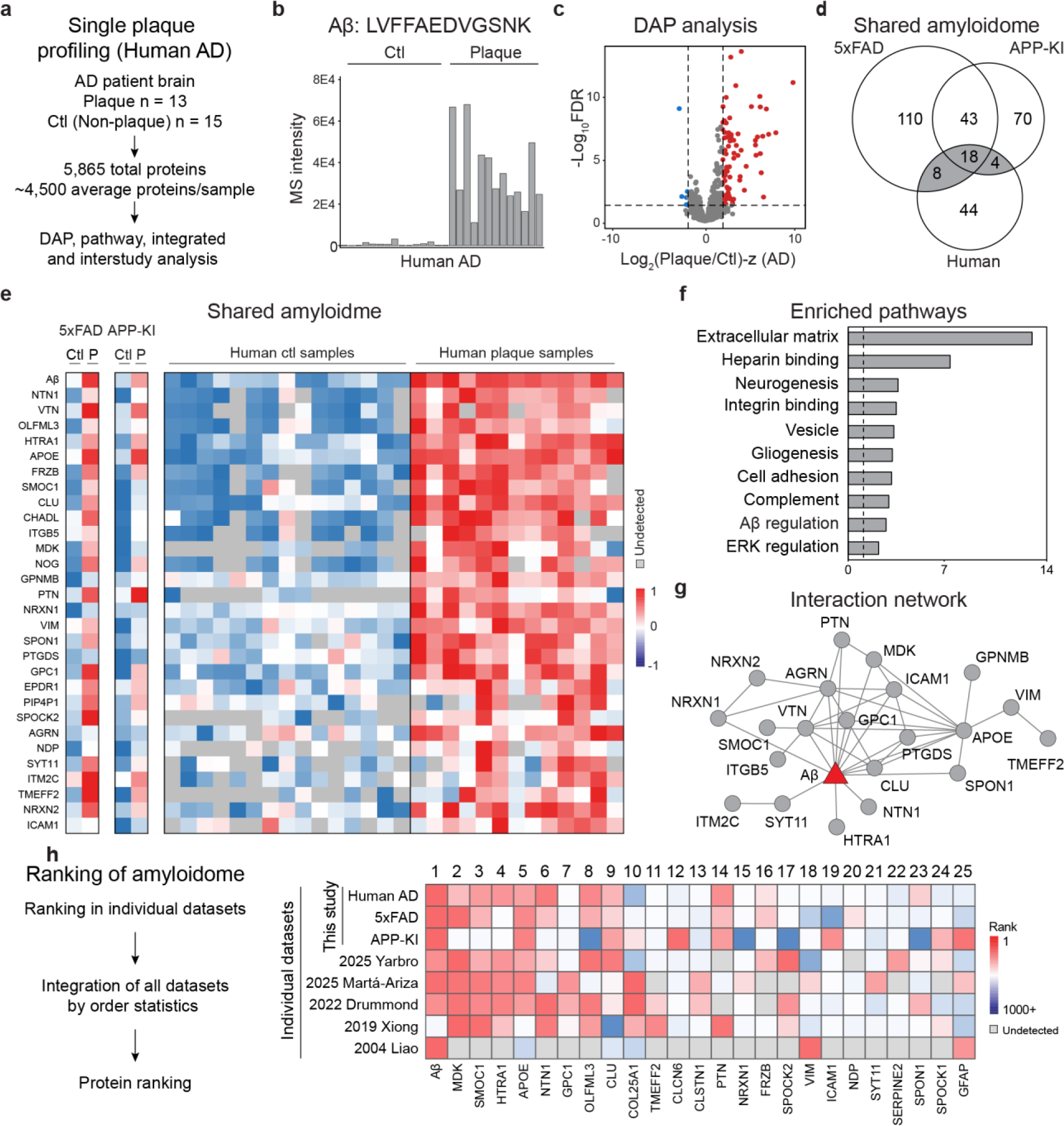
Human plaque proteomics reveals conserved components and identifies core plaque proteins across species. **a**, Paired plaque and non-plaque profiling in human AD (*n* = 28) quantified 5,865 proteins. **b**, MS-based quantification of Aβ using peptide LVFFAEDVGSNK. **c**, Volcano plot of proteins enriched in plaque vs. non-plaque regions. **d**, Overlap of enriched DAPs in mouse and human identifies 38 conserved components. **e**, Heatmap of overlapping DAPs across mouse and human samples. **f**, Pathway analysis highlights extracellular, heparin-binding, and developmental proteins. **g**, PPI network of overlapping DAPs across species. Aβ is highlighted in red. **h**, Enrichment status of overlapping DAPs in prior plaque proteomics studies. **i**, Strategy for interstudy protein ranking using order statistics. **j**, Integrative ranking across eight datasets. The top 25 proteins are displayed, with undetected proteins represented by white boxes.

Differential abundance analysis identified 89 significantly altered proteins (FDR < 0.05; |log₂FC-z| > 2.5), including 84 upregulated in plaques and 5 in non-plaque regions (**Fig. 6c; Table S14**). Among two mouse models and human AD amyloidomes, 30 proteins were consistently enriched between mouse and human, including 18 proteins shared by all three datasets (**Fig. 6d**). The two mouse models shared 73 DAPs, reflecting stronger concordance within mice than across species. A heatmap of shared amyloidome proteins (**Fig. 6e**) revealed a core set of proteins shared across species. Comparisons of these DAPs at each age, models and species are summarized in **Table S15**.

Pathway analysis of DAPs across human and mouse plaques highlighted enriched pathways such as extracellular matrix, heparin binding, neurogenesis, and gliogenesis (**Fig. 6f; Table S16**). Aβ emerged as a central hub protein, connecting with other known components such as APOE and CLU, and less studied proteins such as NTN1, SPON1, MDK, PTN, and NRXN1 (**Fig. 6g**). To determine a core set of amyloidome proteins enriched across amyloid proteomics datasets, we integrated our datasets with five previously published studies (**Fig. 6h**)^17,19,22,36,37^. This cross-study ranking identified the top 25 amyloid-associated proteins, including Aβ, MDK, SMOC1, HTRA1, and APOE (**Table S15**). These findings establish a core list of conserved amyloidome proteins across mouse models and human AD, supported by shared DAPs, overlapping pathways, and hub proteins.

### MDK, PTN, and HTRA1 strongly associated with amyloid plaques in immunostaining

To further validate highly ranked Aβ-associated proteins, we performed immunofluorescence staining in 5xFAD mice (14-month-old) and human AD cases with co-labeling of Aβ peptides (green) with any of Mdk, Ptn, and Htra1 proteins (red). These proteins consistently co-localized with Aβ in the plaque microenvironment, strongly supporting their involvement in the development of amyloid pathology (**Fig. 7a**).

**Fig. 7.**
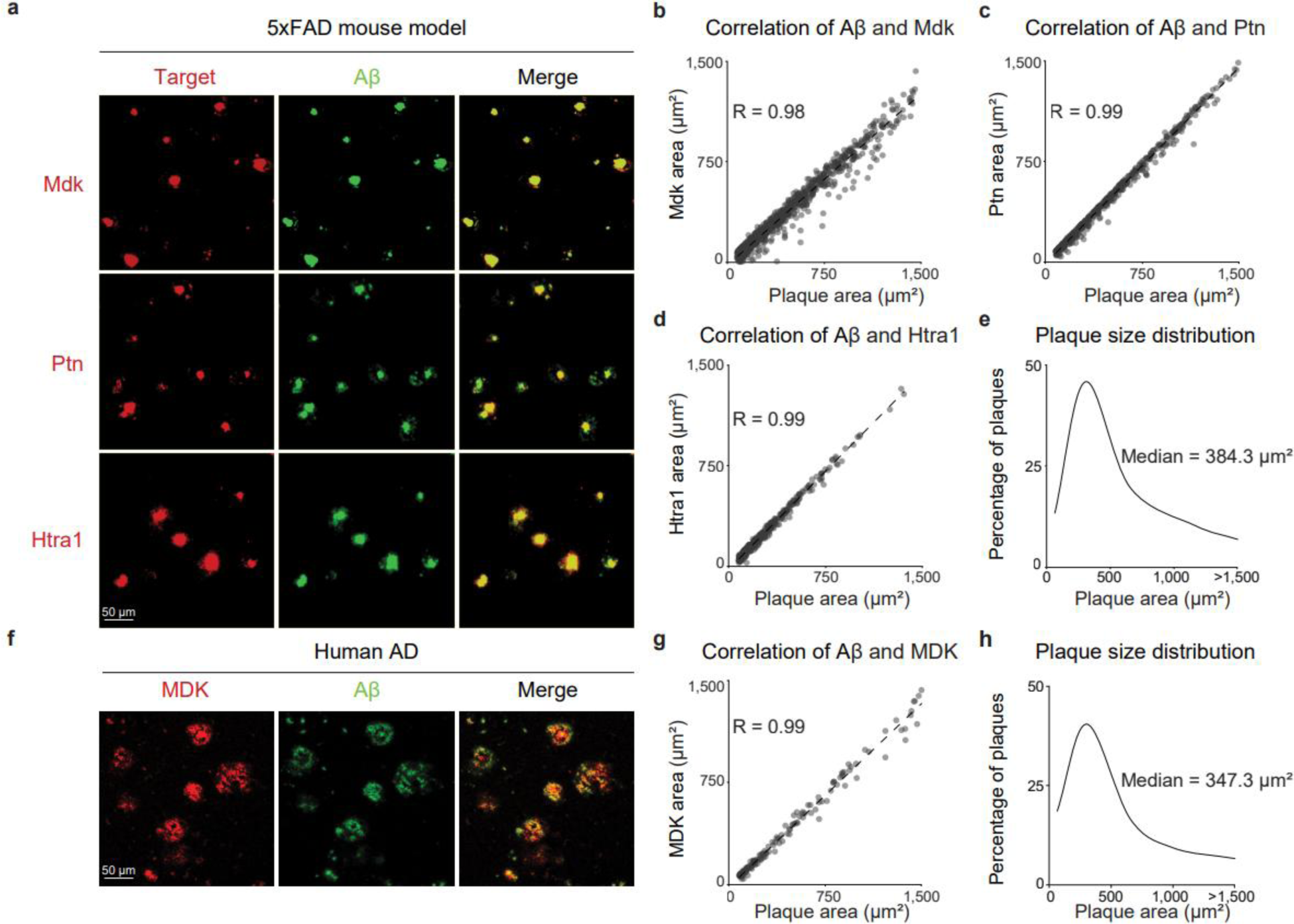
Immunofluorescence validation of MDK, PTN, and HTRA1 across species. **a**, Immunofluorescence labeling of Aβ (green) with Mdk, Ptn, or Htra1 (red) in the cortex of 14M 5xFAD mice. **b-d**, Quantitative correlation analyses between Aβ and Mdk (b), Ptn (c), or Htra1 (d) in 14M 5xFAD mice. **e**, Distribution of plaque size based on Aβ area in 14M 5xFAD mice. f, Immunofluorescence labeling of Aβ (green) and MDK (red) in human AD cortex. **g**, Quantitative correlation analysis between Aβ and MDK in human AD cases. **h**, Distribution of plaque size based on Aβ area in human AD cases.

To extend this to a quantitative level, we analyzed the spatial overlap between Aβ and Mdk, Ptn, or Htra1. In the cortex of 5xFAD mice, 761 plaques were identified and analyzed, yielding a strong correlation between Aβ and Mdk (R = 0.98; **Fig. 7b, Table S17)**. Similar analyses in 501 and 336 plaques showed equally robust correlations for Ptn (R = 0.99) and Htra1 (R = 0.99), respectively (**Fig. 7c; 7d, Table S17**). The distribution of mouse plaque areas exhibited a median of 384.3 µm² and a mean of 557.3 µm² (**Fig. 7e**).

We next examined human AD cases using the same workflow. Co-localization of Aβ and MDK were also evident (**Fig. 7f**), and analysis of 141 plaques confirmed a very strong correlation (R = 0.99; **Fig. 7g**), while the distribution of human plaque areas exhibited a median of 347.3 µm² and a mean of 569.0 µm² (**Fig. 7h**).

Together, these imaging results demonstrate robust co-localization and a strong correlation of Aβ with several core amyloidome proteins in both AD mice and human brain samples.

## Discussion

In this study, we have shown that amyloid plaques form dynamic, temporally ordered molecular niches rather than inert deposits, providing a direct answer to the question of how the plaque microenvironment is composed and remodeled during AD progression. Our single plaque LCM-DIA-MS workflow quantified over 7,000 proteins from ∼50 µm regions of plaque microenvironment and revealed that early plaques are dominated by immune and endolysosomal pathways, whereas later plaques increasingly engage RNA processing and synaptic components. These findings indicate that plaques initiate multicellular responses that change with disease stage. Importantly, these temporal patterns were replicated across two amyloidosis mouse models and supported by conserved signatures observed in human plaques, defining a cross-species core amyloidome that includes APOE, MDK, PTN, and HTRA1. These data suggest that localized proteomic remodeling around plaques provides a mechanistic link between amyloid accumulation and downstream neurodegenerative events.

Our findings extend prior plaque proteomics studies^17,19,22,36,37^ by adding single-plaque resolution and a temporal dimension, offering deeper insight than bulk tissues^13,14^ and detergent-insoluble analyses^38–40^. We observed the same immune, lysosomal, extracellular matrix, and vesicular components consistently enriched in plaque fractions, but with substantially greater depth at the level of individual aggregates. Temporal clustering separated early microglial/astrocytic and lysosomal/vesicular responses from more modest, transient changes involving RNA metabolism and spliceosomal-associated proteins, alongside synaptic alterations. Notably, these RNA-related signals were low at early stages and attenuated at later time points within plaques, suggesting an early adaptive or stress-associated response.

The enriched protein composition further demonstrates an integrated, multicellular niche including reactive microglia, astrocytes, neurons, and extracellular matrix components. As in a previous spatial transcriptomics study that reported localized cellular changes within a radius of ∼20-40 µm of the plaque core^41^, our work focused on a ∼25 µm radius (50 µm dimension) around the Aβ core, capturing associated matrix proteins and the major responding cell types. This revealed a glia-dominated niche; microglia and astrocytes collectively accounted for over half of the plaque microenvironment area. Lysosomal and vesicular enrichment supports sustained glial engagement with aggregated proteins, while synaptic signatures indicate neuronal involvement near plaques. Microglia rapidly respond to early plaque formation^42–44^, and astrocytes exhibit reactive and regionally heterogeneous states with known effects on blood-brain barrier integrity and neuronal signaling^45,46^. Dystrophic neurites surrounding plaques accumulate autophagic vesicles, lysosomal proteins, and hyperphosphorylated tau^11^, emphasizing disrupted proteostasis and axonal transport within the microenvironment. Collectively, these observations define the plaque microenvironment as a glia-dominated but multicellular signaling hub shaped by chronic interactions among glia, dystrophic neurites, and a remodeled extracellular matrix.

Several conserved, Aβ-correlated amyloidome proteins were robustly and consistently enriched across models and species, including MDK, PTN, HTRA1, SMOC1, and others. Many are secreted or extracellular regulators that modulate proteostasis, ECM remodeling, and cell-cell signaling. MDK protein directly binds Aβ, prevents fibril formation, and reduces plaque burden and microglial activation, while loss of MDK worsens pathology^47^. PTN shows similar enrichment and likely participates in overlapping neuroprotective signaling, whereas HTRA1 remodels the extracellular matrix and degrades misfolded Aβ and α-synuclein^48,49^. Their co-enrichment with APOE and complement supports a coordinated network in which plaque-associated growth factors, proteases, and ECM components sustain glial activation and matrix turnover^17^. These findings imply that therapeutic strategies aimed at modulating these multicellular networks may influence plaque-associated pathology.

We acknowledge several limitations in this study. First, the cross-sectional design infers temporal dynamics from age-stratified mice rather than tracking individual plaques longitudinally. Second, although our workflow is optimized for nanogram inputs, sample loss is inevitable and may lead to undersampling^24,50^. Third, we focused on cored plaques; diffuse plaques and vascular amyloid likely contain distinct proteomic signatures^51–53^. Finally, although several proteins were validated by immunofluorescence and cross-species replication, the causal roles of most amyloidome components remain to be determined.

In summary, our quantitative single-plaque proteomics study demonstrates that amyloid plaques are dynamic, conserved pathological microenvironments that coordinate immune-lysosomal responses early and later engage RNA processing and synaptic pathways. These findings advance our understanding of AD pathogenesis by providing molecular resolution of plaque-associated remodeling that was previously masked in bulk analyses. Translationally, conserved plaque-associated proteins and stage-specific pathways offer potential candidates for disease biomarker development and therapeutic targeting.

## Methods

### AD mouse models and human tissues

All mice were maintained on a C57BL/6J background (Jackson Laboratory #000664). This study included 5xFAD mice (Jackson Laboratory #034848-JAX)^25^ and APP^NLGF/NLGF^ mice^26^. Mice were sacrificed, and their brains were dissected, washed with phosphate buffer solution (PBS), embedded in O.C.T. (optimal cutting temperature compound), and stored at −80 °C. Three to four 15 μm thick cryosections were obtained on a CryoStar NX 70 (Thermo Fisher Scientific) and placed on charged glass slides (Superfrost Plus, Thermo Fisher Scientific) for plaque visualization, or on PEN Membrane Glass Slides (Thermo Fisher Scientific) for LCM.

Human postmortem brain tissue samples (frontal cortex) were provided by Banner Sun Health Research Institute^54^. A total of five brain tissue samples (**Table S1**) were used for single plaque proteomics profiling.

### Plaque visualization and laser capture microdissection

Cryosections (12 µm thickness) were equilibrated at 21 °C, fixed for 5 min in 75% ethanol, and stained with X-34 (10 µM) for 2 min. The sections were then washed sequentially with PBS, 75% ethanol, 95% ethanol, and twice with 100% ethanol. Finally, sections were air-dried for 5 min or stored at −20 °C. LCM was performed by automatic laser pressure catapulting (AutoLPC) on the Accuva Cellect system (Laxco), with the following settings: 10x magnification; energy, 70; focus, 78 (both in arbitrary units); 17 µm distance between AutoLPC shots; 3 µm distance from the laser line, and diagonal AutoLPC shooting pattern. Fluorescently labeled plaques were microdissected from various regions within the cortex for each case, with paired non-plaque regions from adjacent areas as controls for local proteomic background. The diameter of each collection was typically ∼50 µm onto CapSure Macro LCM Caps (Thermo Fisher Scientific) and stored at −80 °C.

### Sample preparation for LC-MS/MS

Samples (one capture) were incubated with 2 μL of lysis buffer (100 mM ammonium bicarbonate (ABC), 5 mM dithiothreitol (DTT), 0.02% n-dodecyl β-D-maltoside (DDM), 10 ng/μL trypsin, pH 8.0) at 47 °C for 30 min. Samples were further digested and alkylated by adding 0.5 μL of digestion buffer (100 mM ABC, 0.02% DDM, 50 ng/μL trypsin, pH 8.0) and incubated at 37 °C for 30 min. Samples were acidified by adding 1 μL of 20% formic acid (FA) after centrifuging at 6,000 x g to collect all solutions in a 0.5 mL low-binding tube. The solution was clarified by centrifugation at 21,000 x g for 10 min and transferred to inserts for LC loading.

### Label-free DIA MS

Peptides were analyzed with optimized settings^30^, using a C18 column (10 cm x 50 µm, 1.9 µm particle size; 55 °C; Bruker Daltonics) on a nanoElute 2 liquid chromatography system (Bruker Daltonics). A 45-minute gradient was applied at a flow rate of 0.25 µL/min, increasing buffer B (0.1% formic acid in acetonitrile; buffer A: 0.1% formic acid in water) from 5% to 18%, followed by a ramp to 65% for post-run column washing.

MS data were acquired using the diaPASEF method on a timsTOF SCP mass spectrometer (Bruker Daltonics). The capillary voltage was set to 1500 V. The MS1 and MS2 spectra were acquired from 100 to 1700 m/z with an ion mobility range of 0.7 to 1.6 Vs/cm^2^. The accumulation and ramp time were set as 166 ms. Collision energy was applied using a linear ramp from 20 eV at 1/K0 = 0.6 Vs/cm^2^ to 59 eV at 1/K0 = 1.6 Vs/cm^2^. Isolation windows of a 20 m/z width were used to cover the mass range of 400 to 1200 m/z.

### Protein identification and quantification

The raw files (. d file) from timsTOF were analyzed in DIA-NN (version 1.8) with the library-free mode, using an in silico spectral library and match between run (MBR). Protein FASTA files from UniProt database (49,775 and 82,156 entries for mouse and human) were used for library generation with the following settings: precursor FDR 1%; mass accuracy at MS1 and MS2 as 0 (automatic); scan window set to 0 (automatic); trypsin/P with maximum 2 missed cleavages; carbamidomethylation on Cys as fixed modification; oxidation on Met as variable modification; variable maximum modifications set to 2; peptide length from 7 to 30; precursor charge 1-4; precursor m/z from 300 to 1,800; fragment m/z from 200 to 1,800. The search results were further filtered with *q* value < 0.01 for precursor and protein groups at the library level. Cross-run normalization was performed after quantification based on the median protein intensity and loading amount of each sample if necessary.

### Data preprocessing and differential abundance analysis

The following steps were implemented for quality control. (i) The protein expression matrix was filtered to remove common contaminants (e.g., trypsin and keratin) and proteins detected in fewer than three samples. (ii) Aβ tryptic peptides were extracted as a positive control to evaluate the quality of the LCM captures. (iii) For the accepted proteins with missing values, we first used the number of detected samples as a proxy for quantification and compared plaque versus control samples using the G test^55^. If the resulting p value was < 0.05, missing values were imputed using the sample-specific minimum value and adjusted to ensure an equal number of detections across two groups. We did not fill all missing values to avoid over-interpretation. (iv) Paired moderate *t* tests were performed to compare protein abundance between plaques and the non-plaque area, followed by the Benjamini–Hochberg method to generate FDR values. (v) We filtered the significant differentially abundant proteins (DAPs) based on FDR <0.05, and absolute z values of log2 fold change (log2FC-z): > 2.5 in mouse and > 2 in human. Data from each age were analyzed separately and then normalized within 5xFAD, APP-KI, and human.

### Temporal clustering

Temporal profiling of plaque-associated proteins was performed in mouse samples to assess dynamic changes during disease progression. Log_2_-transformed DAP intensity values from plaque samples were used as input. For each protein, mean abundance was calculated within each age group, then normalized by subtracting the 3-month mean (log scale) to obtain relative fold changes across ages. Proteins showing opposite fold-change directions between 8- and 14-month comparisons were excluded. Remaining proteins were clustered based on temporal fold-change trajectories, guided by WGCNA results and manual inspection^56^.

### Multiscale co-expression network analysis using MEGENA

Multiscale Embedded Gene Co-expression Network Analysis (MEGENA) was performed to construct multiscale co-expression networks from normalized protein abundance data. Prior to analysis, missing values were filtered to retain proteins with less than 50% missingness across samples. Protein abundance values were median-centered and merged across conditions for downstream network inference. Pairwise Pearson correlations were computed among all proteins, and significance was assessed using permutation-based FDR < 0.05. Significant correlations were used to construct a Planar Filtered Network (PFN) and multiscale clustering analysis (MCA) was then applied to the PFN to identify modules of co-expressed proteins using a module significance threshold of p < 0.05 and hub connectivity significance p < 0.05 based on random tetrahedral networks. Modules with fewer than 10 nodes or exceeding half the network size were excluded. Hub proteins were defined as nodes with statistically significant connectivity within each module. Module enriched in DAPs were performed using fgsea package with 10,000 permutations and ranked gene statistics derived from log₂FC-z for each comparison (5xFAD and APP-KI). All analyses were performed in R (v4.4.0) using the MEGENA package with parallel computation (40 cores).

To identify biological modules associated with plaque pathology, we performed module-trait correlation analysis following the framework of the Weighted Gene Co-expression Network Analysis pipeline. Module eigengenes were computed to represent the overall expression profile of each co-expression cluster. The eigengene matrix was then correlated with binary phenotypic traits corresponding to plaque and non-plaque regions across AD mouse models. Pearson correlation coefficients and corresponding p values were computed for each module-trait pair, and statistical significance was determined using an FDR threshold of 0.05 and an absolute correlation coefficient cutoff of 0.4.

Module annotations for each MEGENA module were performed using the clusterProfiler package. Briefly, module membership data were imported, and unique modules were iteratively processed to test for GO Biological Process enrichment with the org.Mm.eg.db database. For each module, the top ten enriched GO terms were identified based on adjusted P value ranking. These terms were then qualitatively assessed to derive the module annotation.

### Pathway enrichment and protein-protein interaction network analysis

Pathway enrichment of all DAP sets was performed by Gene Ontology (GO) enrichment and further analyzed by the PANTHER overrepresentation test (Fisher’s Exact test)^57,58^. Pathway enrichment was annotated with emphasis to GO categories relevant to neurodegeneration, informed by prior knowledge of amyloid pathology and AD pathogenesis. Results were filtered (FDR < 0.01, enrichment > 2) to identify highly confident pathways. To construct PPI networks, DAP sets were used as input to STRING using mouse or human as organism with default settings^59^.

### Prioritization of amyloidome proteins based on amyloid plaque proteomics datasets

To identify proteins consistently colocalized with plaques and potentially involved in plaque dysfunction, we applied an order statistic-based ranking method for multi-dataset integration^14^. This approach combines 𝑁 independent protein rankings into a single consensus list. Eight amyloid plaque proteomics datasets were integrated, including the human and mouse datasets from this study and five previously published studies^17,19,22,36,37^. Within each dataset, proteins were ranked by p- or q-value, or by a combined measure of significance and fold change, depending on the available data.

### Other Bioinformatics Analysis

Most data visualization was performed in R and Excel. The following R packages were used to visualize the data: ‘ggplot2’ (boxplots, PCA plots, volcano plots, correlation plots and column chart), ‘pheatmap’ (heatmaps), ‘ggVennDiagram’ (Venn diagram) and ‘UpsetR’ (UpSet plots). **Fig. 1b** was generated with the assistance of BioRender.

### Immunostaining imaging

For Mdk, Ptn and Htra1, staining procedures followed a protocol adapted from Bai et al^55^. Primary antibodies included Mdk (1:100, R&D Systems, Cat# AF-7769), Ptn (1:500, Abcam, Cat#79411), Htra1 (1:100, Abclonal, NP_062510.2) and Aβ (1:100, Immuno-Biological Laboratories, Cat# 10323). Sections were incubated with primary antibodies overnight at 4 °C. For Aβ or Ptn, sections were incubated fluorophore-conjugated secondary antibodies (1:500, Jackson ImmunoResearch Laboratories). For Mdk or Htra1 staining, HRP polymer-conjugated secondary antibodies (Vector Laboratories) were applied, followed by detection with Cyanine 5 tyramide. Slides were mounted with Aqua-poly/Mount (PolySciences, Cat#18606).

Similarly, for the cell type marker staining, of IBA1 (1:1000, BioCare, CP 290 A), GFAP (1:5000, Dako, Z0334), MAP2 (1:500, Sigma Aldrich, AB5622), NEUN (1:1000, Cell Signaling Technology, Cat#24307), and Biotin Aβ (1:1000, Immuno-Biological Laboratories, Cat#10326) were applied as part of a multiplex immunofluorescence panel. Discovery QD DAPI nuclear counterstain was applied, and the slides were mounted with Prolong Gold Antifade reagent containing DAPI (Thermo Fisher Scientific, P36935). The stained slides were scanned using a Zeiss AxioScan Z1 slide scanner (Carl Zeiss Microscopy, LLC). Images were acquired using a 20X/0.8 NA objective.

### Quantification of distinct cells in the plaque microenvironment

Immunohistochemistry staining was performed, and whole-slide images were acquired using a Zeiss Axioscan.Z1 scanner. Circular regions of interest (diameter = 50 μm), matching the size used for LCM capture, were placed on Aβ-positive plaque areas. Within each region, the *cell detection* function in QuPath^32^, an open-source digital pathology platform, was applied to quantify the positive areas for cell-type markers including Iba1, Gfap, NeuN, and Map2. The detection threshold was set at one-third of the channel minimum, with cell expansion parameters ranging from 0 to 1.5 μm. The positive areas were exported and used to calculate the percentage of each cell marker area within the amyloid plaque region. Aβ, microglia, astrocytes, and neurons were quantified using antibodies against Aβ, Iba1, Gfap, and the sum of NeuN and MAP2, respectively.

### Quantification of proteins in plaque immunohistochemistry

Quantitative analysis was conducted with QuPath. Automatic plaque detection was achieved using a trained pixel classifier for Aβ within the selected cortical regions. Within the Aβ-positive areas, additional trained classifiers for Mdk, Ptn, and Htra1 were applied to delineate the corresponding protein-positive regions on the same slides. The measured areas of Aβ, Mdk, Ptn, and Htra1 were extracted, and correlation coefficients were calculated after excluding plaques with areas exceeding 1,500 μm² or smaller than 80 μm².

## Data availability

The mass spectrometry proteomics data have been deposited to the ProteomeXchange Consortium via the PRIDE partner repository with the dataset identifier PXD061248 and PXD061291.

## Acknowledgements

We thank Dr. Takaomi Saido for providing the APP-KI mice. We also thank the St. Jude Shared Resources and Core Facilities, including Proteomics and Metabolomics Center, Animal Research Center, and Cell and Tissue Imaging Center. This work was partially supported by National Institutes of Health grants R01AG092468, RF1AG064909, RF1AG068581, R01AG053987, U19AG069701, and the ALSAC foundation. The Banner Sun Health Research Institute Brain and Body Donation Program was supported by National Institutes of Health grants U24NS072026, P30AG072980, P30AG019610, the Arizona Department of Health Services, the Arizona Biomedical Research Commission, and the Michael J. Fox Foundation for Parkinson’s Research.

## Author contribution

J.P. and Y.J. conceived the project. J.P., G.Y., and T.G.B. acquired funding. M.C., J.W., Y.J., P.- C.C, H. Sun, M.N., Z.W., S.H., Z.W., H.T., A.A.H., M.L., H. Sheppard, and J.P. performed the experiments. G.E.S., T.G.B., M.D., and D.W.D. characterized and provided human brain samples. J.M.Y., J.W., M.C., H.K.S., Y.F., Z.F.Y., A.Z., X.W., and J.P. analyzed the data. J.W., J.M.Y., M.C., and J.P. wrote the manuscript.

## Additional information

Extended data is available for this paper.

## Extended Data Figures

**Extended Data Fig. 1.**
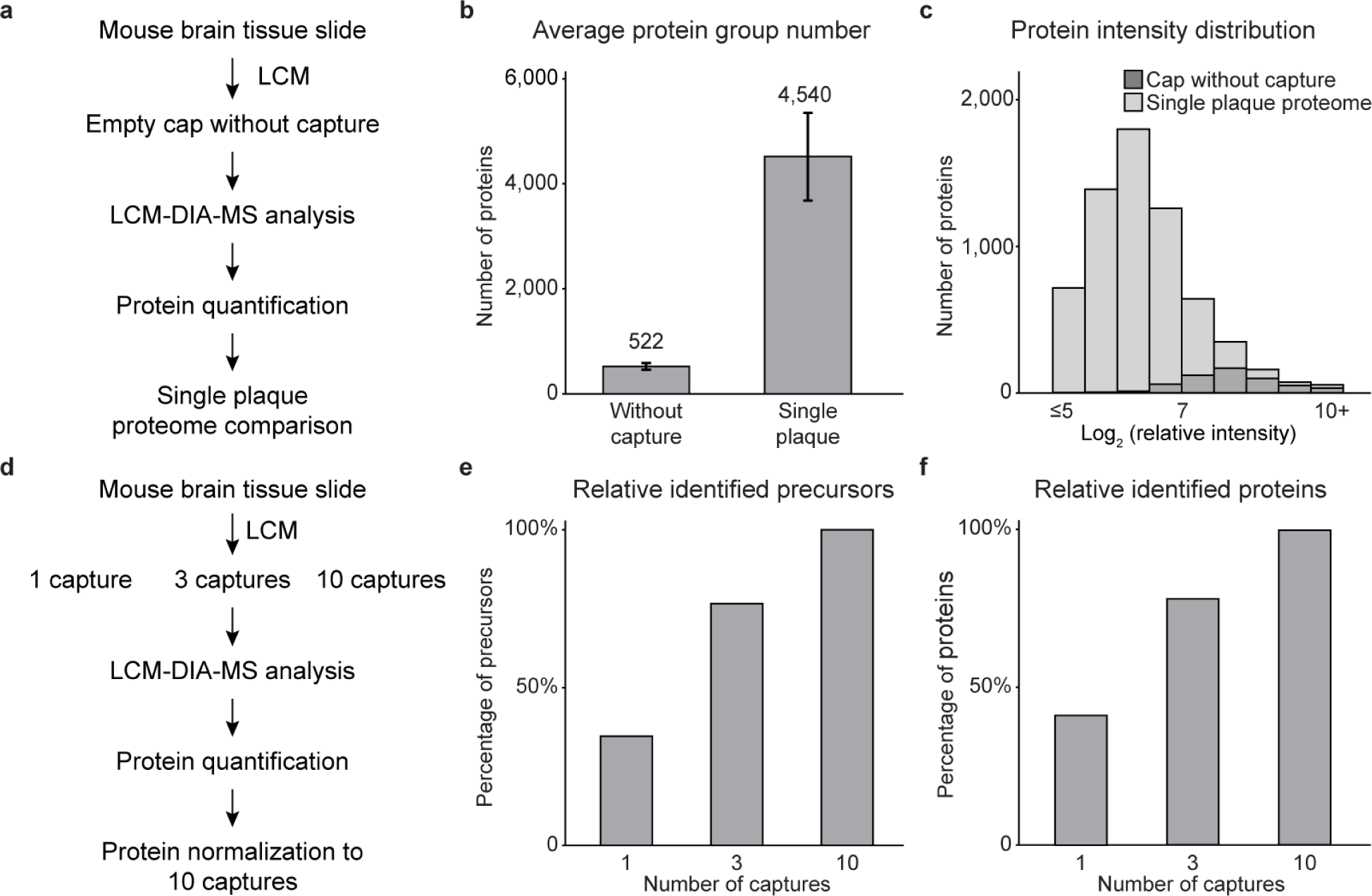
Evaluation of the LCM-DIA-MS pipeline. **a**, Experimental design for comparing single-plaque and negative-control samples using the LCM-DIA-MS workflow. **b**, Average identified proteins of single-plaque and negative-control LCM samples. **c**, Distribution of protein intensities from single-plaque and negative-control samples. **d**, Design of pipeline evaluation using LCM capture titration. **e, f**, Numbers of identified precursors and proteins from 1, 3, and 10 LCM captures.

**Extended Data Fig. 2.**
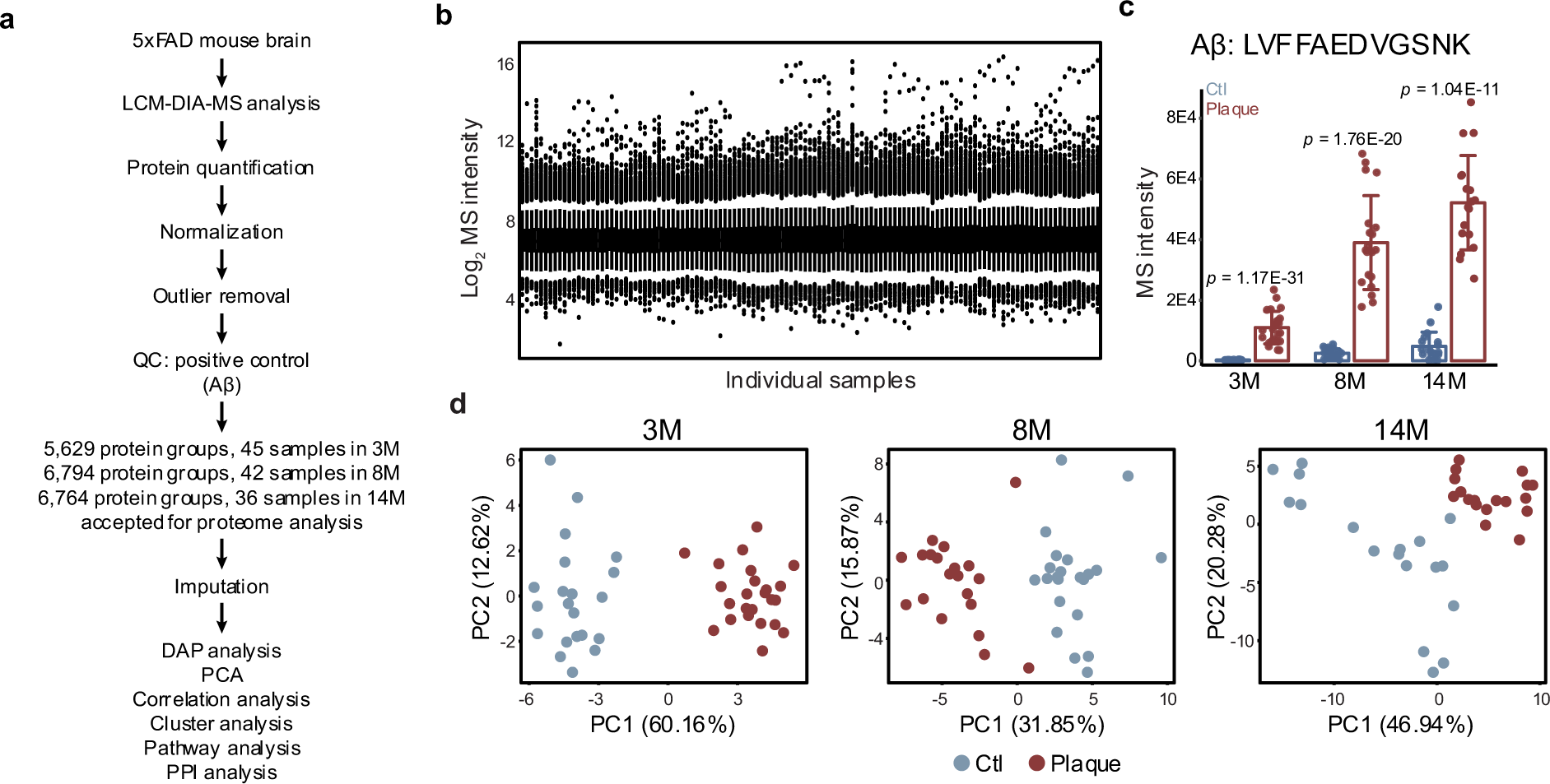
Single-plaque proteomic profiling across disease progression in 5xFAD mice. **a**, Overview of quality control and data processing workflow for the 5xFAD model. **b**, Distribution of normalized protein intensities across all LCM samples. **c**, Quantification of Aβ peptide (LVFFAEDVGSNK) intensities in plaque and non-plaque regions across three disease stages. **d**, PCA distinguishing plaque and non-plaque proteomes at each age.

**Extended Data Fig. 3.**
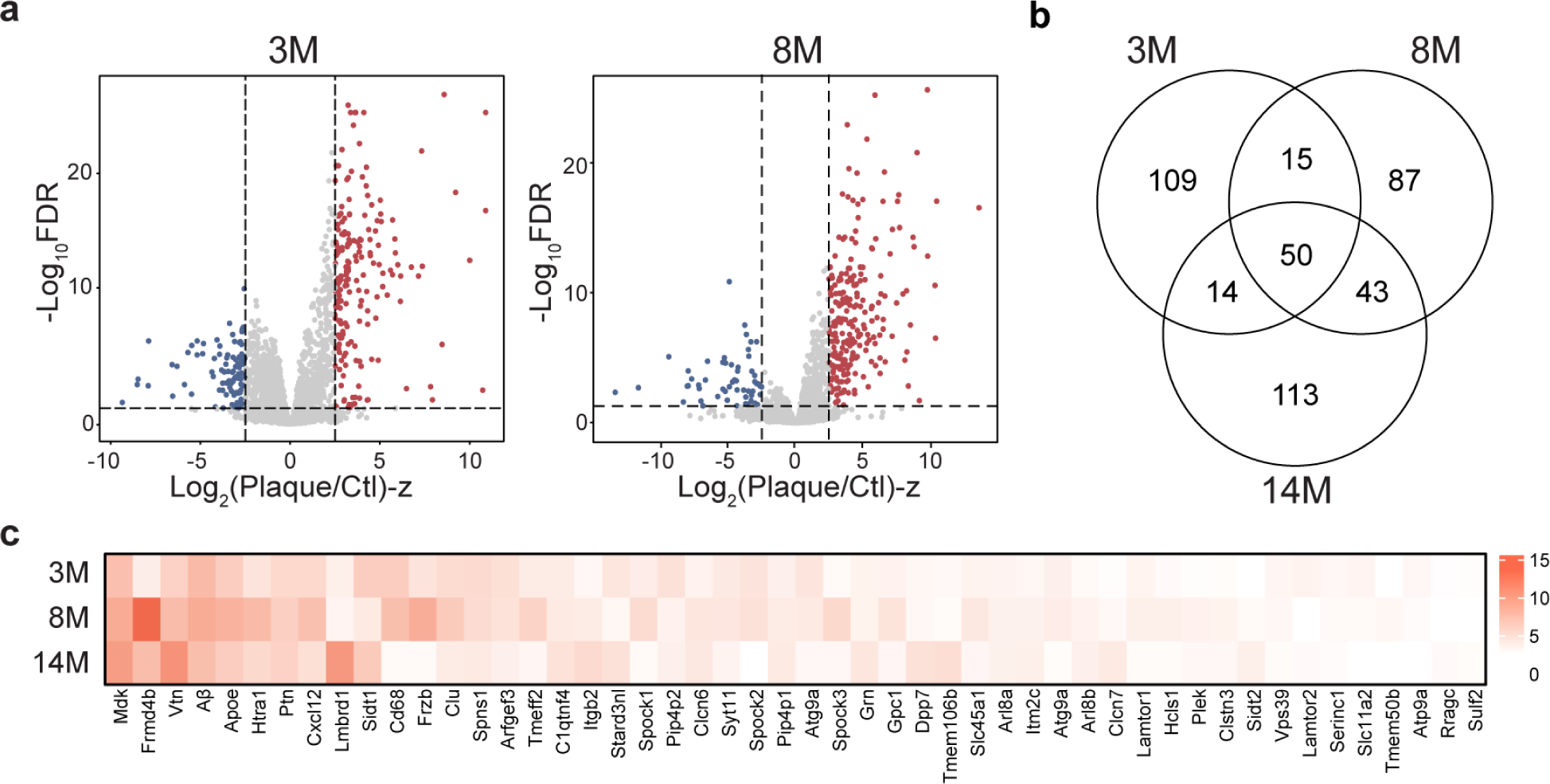
DAP analysis across three ages in 5xFAD mice. **a**, DAP analysis comparing plaque and non-plaque regions at 3M and 8M. **b**, Venn diagram illustrating the overlap of DAPs identified at the three ages. **c**, Heatmap showing the Log₂FC-z values of the overlapping upregulated DAPs across all ages.

**Extended Data Fig. 4.**
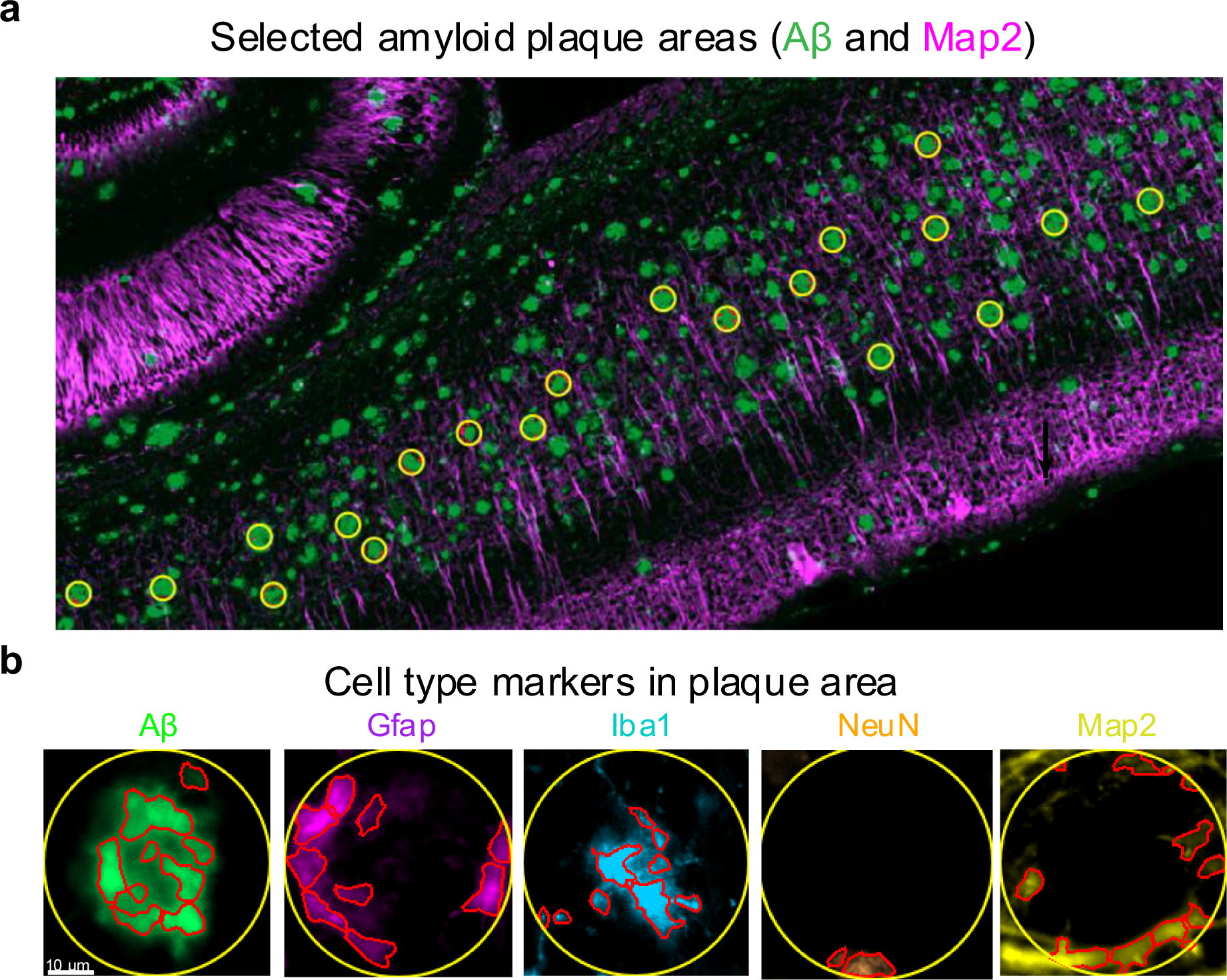
Cell-type detection in the amyloid plaque microenvironment of 14-month 5xFAD mice. **a**, Representative image showing selection of amyloid plaque microenvironments in the cortical region. **b**, Immunofluorescence labeling of cell-type markers within the selected amyloid plaque microenvironments.

**Extended Data Fig. 5.**
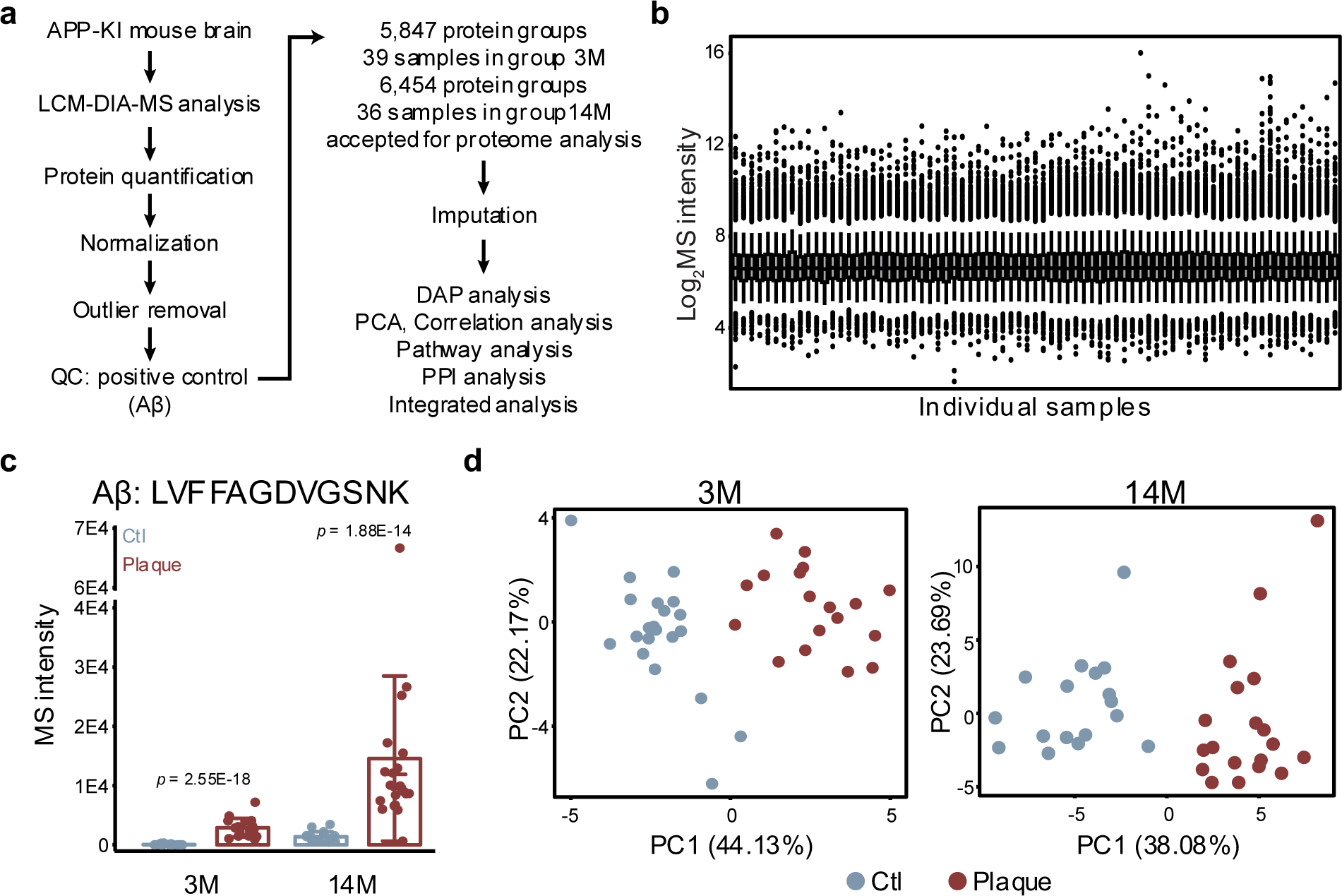
Single-plaque proteomic profiling across disease progression in APP-KI mice. **a**, Overview of quality control and data processing workflow for the APP-KI model. **b**, Distribution of normalized protein intensities across all LCM samples. **c**, Quantification of Aβ peptide (LVFFAGDVGSNK) intensities in plaque and non-plaque regions across three disease stages. **d**, PCA distinguishing plaque and non-plaque proteomes at each age. **e**, DAP analysis between plaque and non-plaque samples at 3M. **f**, Integrated analysis of overlapping DAPs across different time points and models.

**Extended Data Fig. 6.**
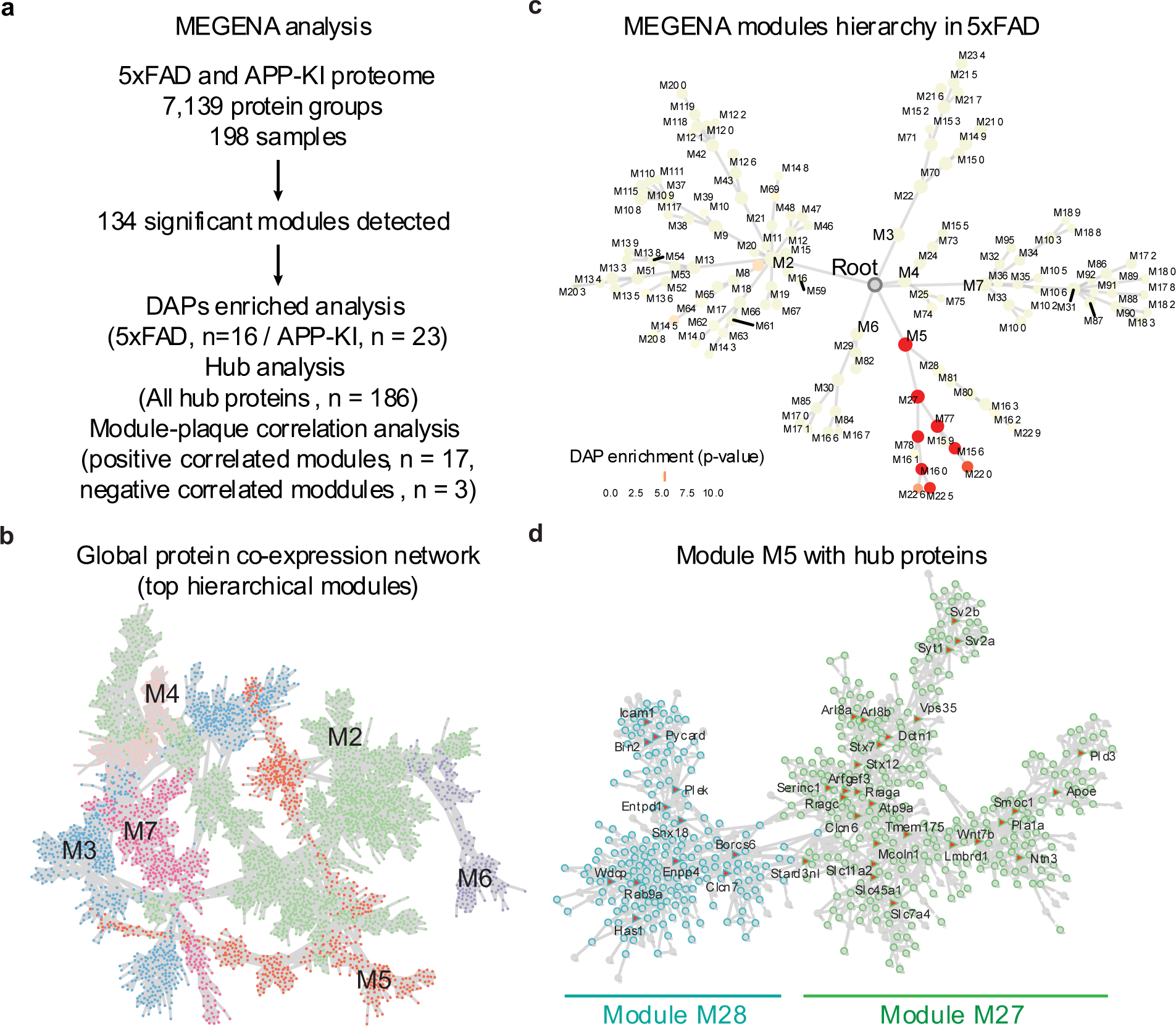
Modules identified from MEGENA. **a**, Overview of MEGENA across all 5xFAD and APP-KI samples. **b**, Global protein co-expression network in the top hierarchical modules. **c**, MEGENA modules hierarchy in 5xFAD showing the enrichment of DAPs. **d**, The network and hub proteins in Module 5.

**Extended Data Fig. 7.**
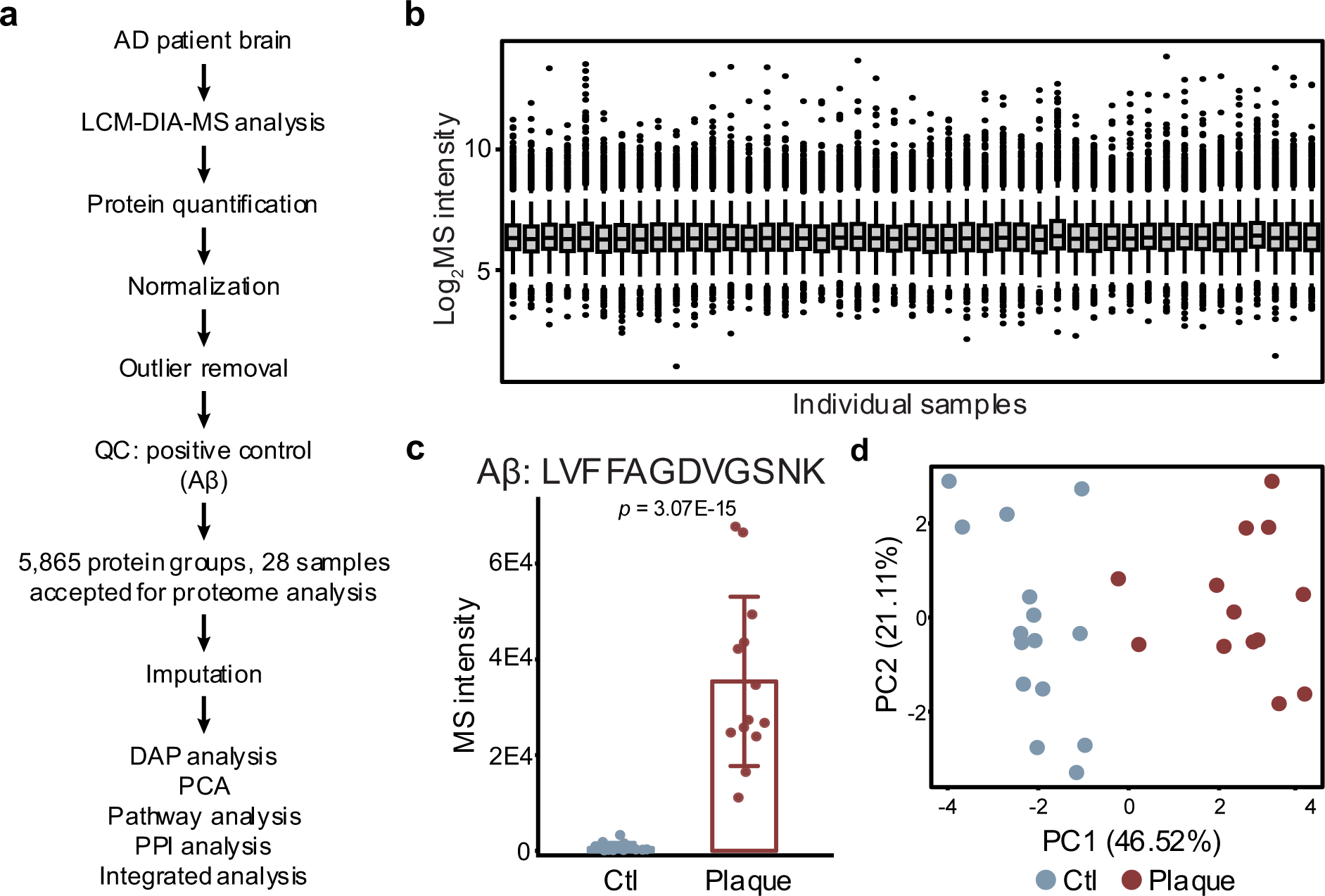
Single-plaque proteomic profiling across disease progression in human AD cases. **a**, Overview of quality control and data processing workflow for the human AD cases. **b**, Distribution of normalized protein intensities across all LCM samples. **c**, Quantification of Aβ peptide (LVFFAGDVGSNK) intensities in plaque and non-plaque regions. **d**, PCA distinguishing plaque and non-plaque proteomes. **e**, Integrated analysis of overlapping DAPs across different time points, models and species.

